## Supplementary Figure 1 for "Multiscale virtual screening optimization for shotgun drug repurposing using the CANDO platform"

August 2020

Figure 1: **Comparison of drug-indication pair rankings for two CANDO platform pipelines.** For each pipeline, 13,746 drug indication pairs are ranked relative to each other along each axis. Points (which may overlap) denote a drug-indication pair plotted according to its corresponding rank value in each pipeline. The panels depict the total distribution of rankings, as well as different subsets, at linear and log scales. Panel (a) depicts the ranking of all drug-indication pairs, and (b) is the same distribution plotted on a logarithmic scale. (c) depicts two subsets plotted on linear and logarithmic scales of the drug-indication ranking distribution of each pipeline containing highly ranked drug-indication pairs (each drug-indication pair ranked higher than 373 and 100 for each pipeline). (d) depicts subsets of the ranking distribution where each pipeline ranked a drug substantially highly in at least one pipeline but not necessarily the other (i.e. higher than 100 for one pipeline and any rank for the second pipeline). The highest density region of the drug-indication pair rankings distribution indicates pairs that are ranked relatively high by each pipeline. There is substantial consensus between each pipeline in terms of ranking pairs relatively highly. However, the asymmetric distribution of many pairs (particularly those that are ranked at a high accuracy threshold for only one pipeline) suggests some differentiation on a per drug-indication pair basis, giving rise to per indication differences and contributing to the enhanced performance of the hybrid decision pipeline.

(a) **Ranking of all drug-indication pairs for two pipelines.**

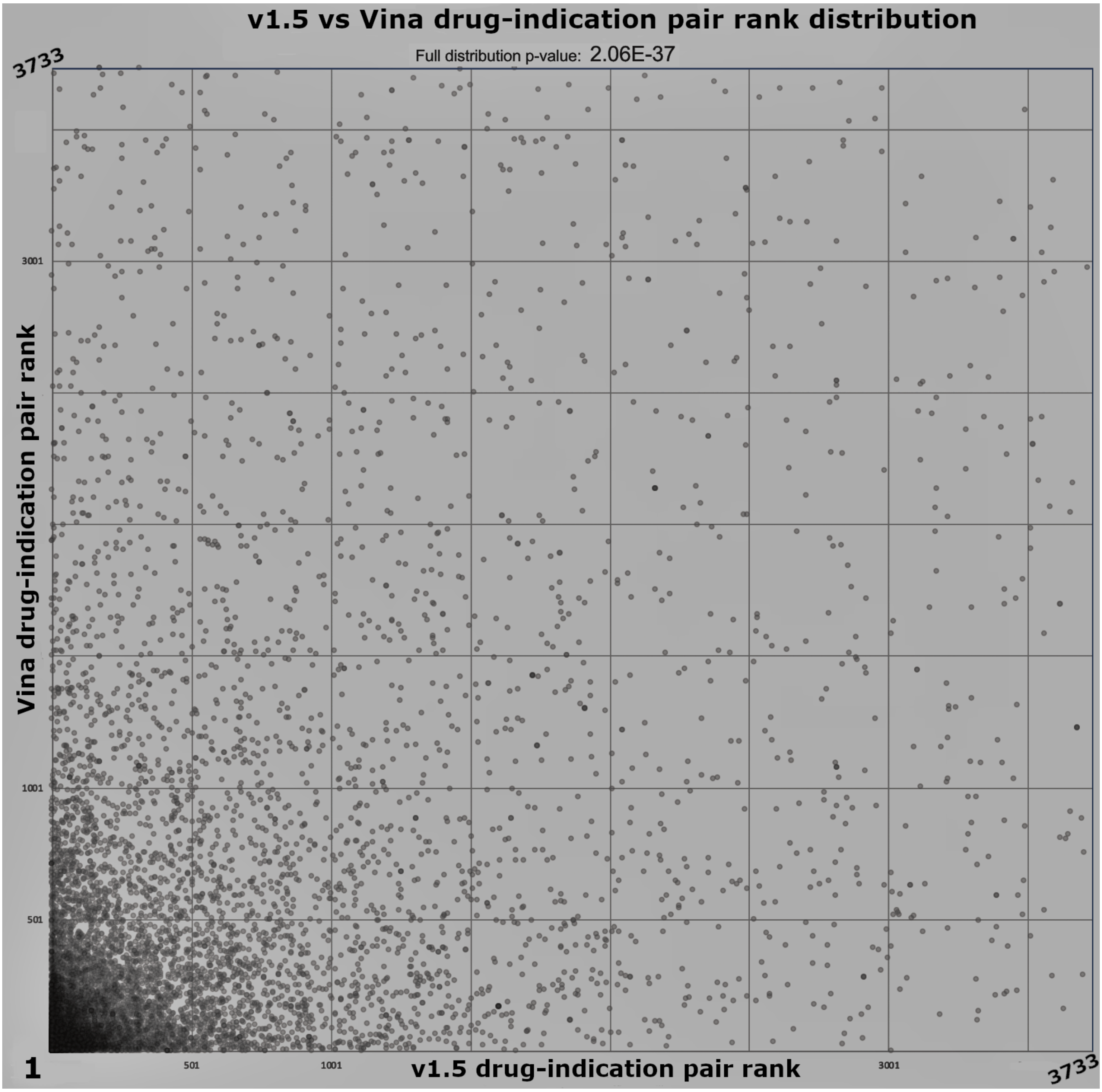

(b) **Drug-indication ranks for two pipelines on a logarithmic scale.**

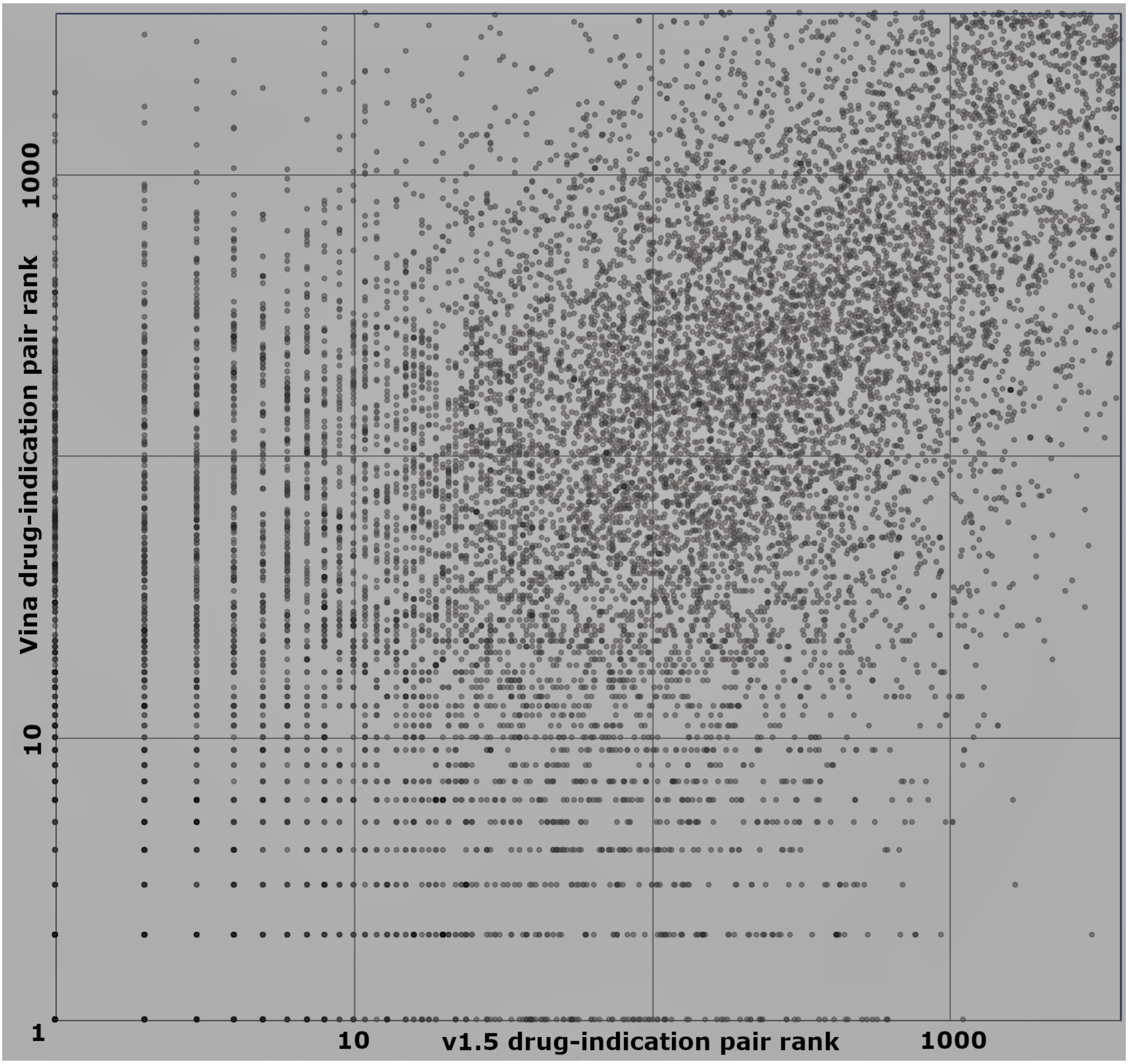

(c) **High ranking drug-indication pair subset distribution.**

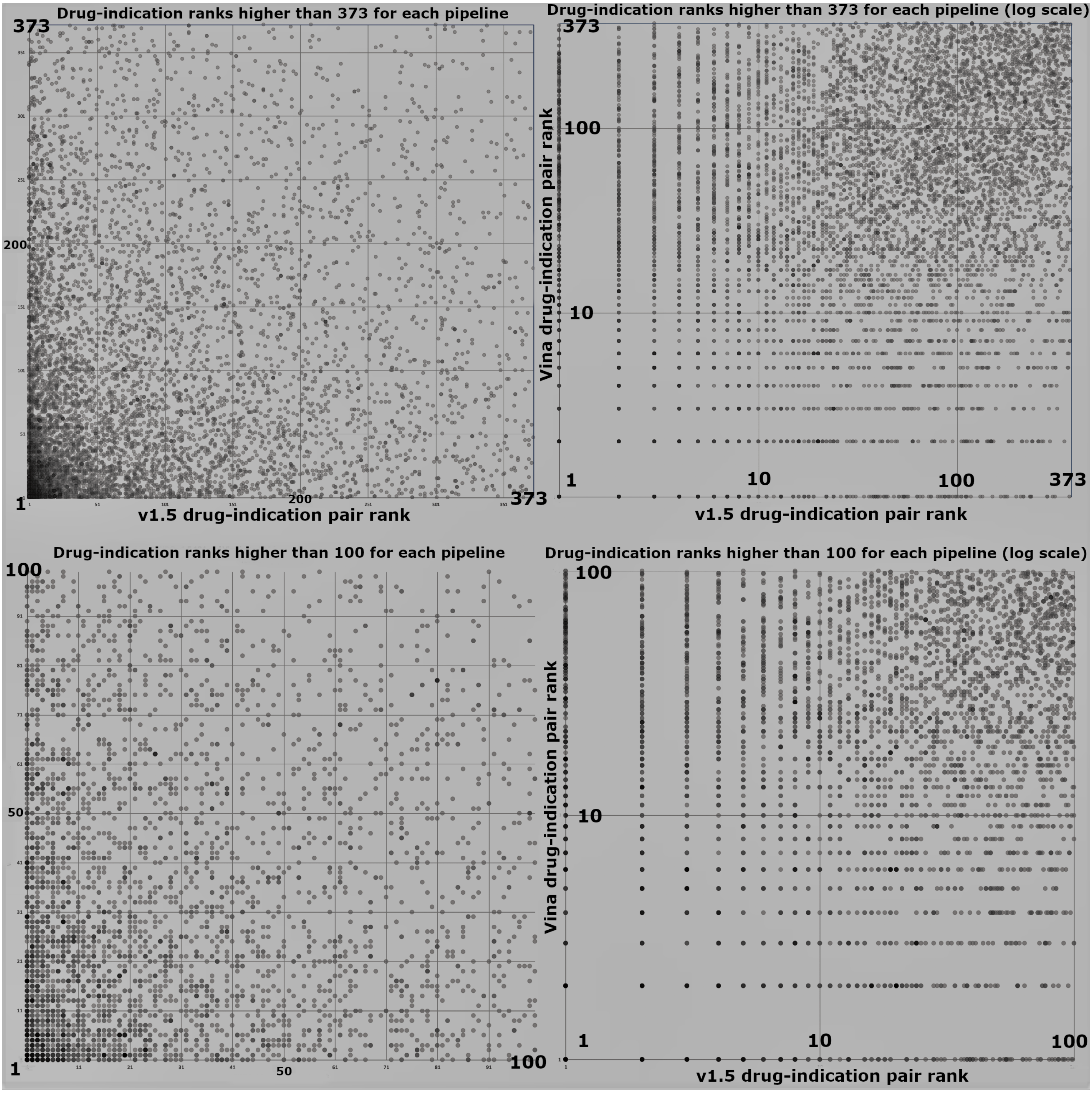

(d) **Subset of drug-indication pairs ranked highly by one pipeline, but not necessarily by the other.**

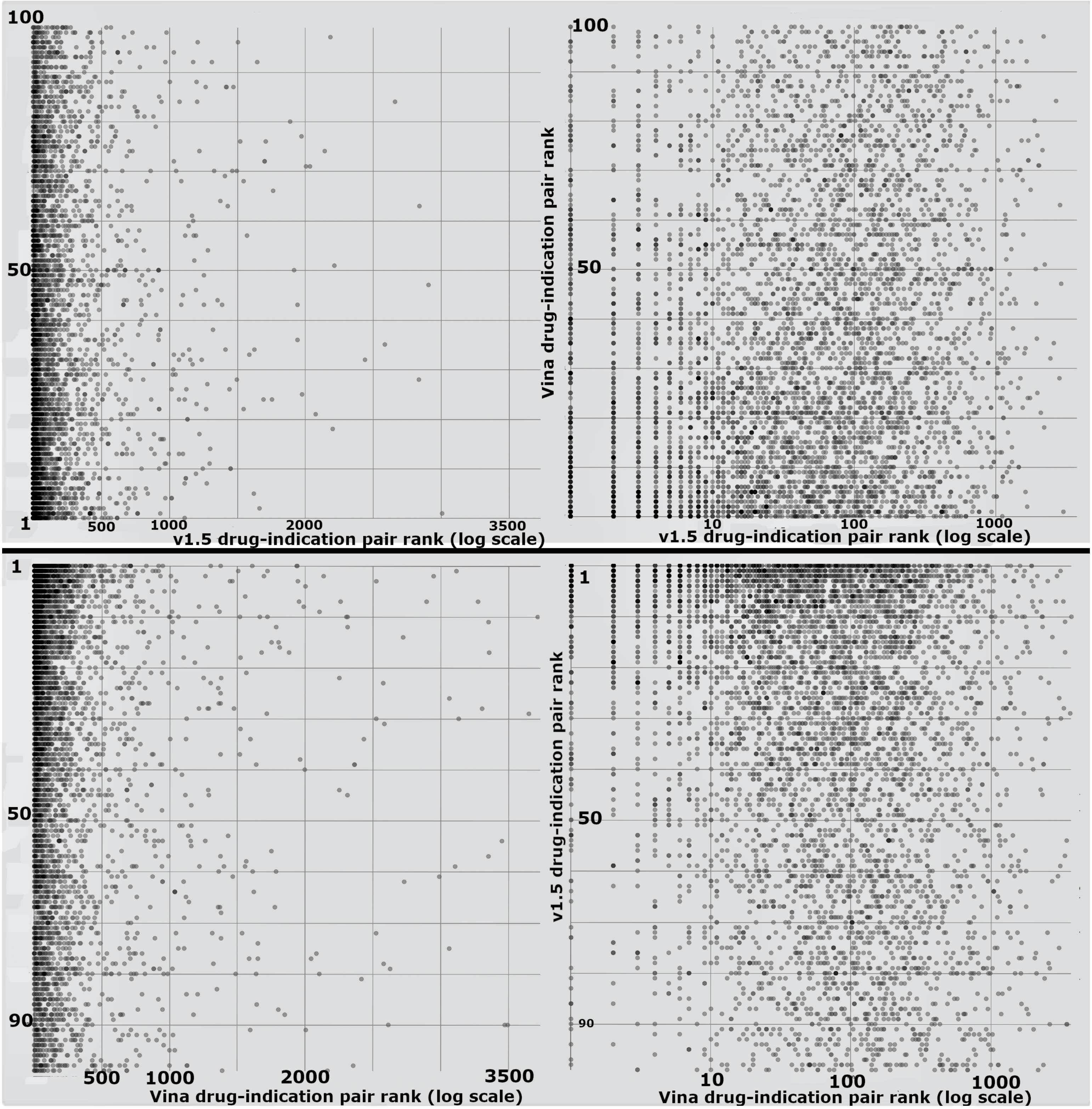
